# A novel tripod probe and lateral flow test to improve CRISPR/Cas12a assay: benefits of branched probe based on trebler phosphoramidite modification

**DOI:** 10.1101/2025.05.29.656603

**Authors:** Irina V. Safenkova, Maria V. Kamionskaya, Aleksandr V. Ivanov, Anatoly V. Zherdev, Boris B. Dzantiev

## Abstract

CRISPR/Cas12a-based assays, when integrated with lateral flow tests (LFTs), provide highly specific nucleic acid detection in a simple, rapid, and equipment-free format. Nevertheless, traditional DNA probes utilized for cleavage by Cas12a have notable limitations as the cleaved probe only has one label. To overcome this challenge, we engineered a novel type of DNA probe with multiple fluorescein (FAM) labels and a biotin-labeled single-stranded DNA fragment (polyFAM probe). The cleaved polyFAM parts of probes were detected using a specially designed sandwich LFT, where FAM-specific antibodies were immobilized in the test zone and conjugated with gold nanoparticles. The LFT ensured accurate recognition of the cleaved polyFAM fragments within 10 minutes. A comparison of five distinct polyFAM probes revealed that the highest signal-to-noise ratio was achieved with a tripod-branched probe synthesized via trebler phosphoramidite modification. Each arm of the tripod probe consists of a hexaethylene glycol spacer ending in a FAM label. Upon Cas12a cleavage, the tripod structure carrying three FAMs is released and detected by LFT. A rapid magnetic separation strategy was subsequently implemented, facilitating the efficient removal of uncleaved probes via biotin–streptavidin capture within 5 minutes. The CRISPR/Cas12a–tripod–LFT strategy demonstrated excellent sensitivity without preamplification, with a detection limit of 1.4 pM for DNA target of *Salmonella* Typhimurium. The CRISPR/Cas12a-tripod-LFT with preliminary loop-mediated isothermal amplification enabled the detection of as few as 0.3 cells per reaction. This innovative tripod probe with corresponding LFT creates a universal, sensitive, rapid, and equipment-free biosensing platform for CRISPR/Cas12a-based diagnostics in point-of-care applications.

## INTRODUCTION

Point-of-care testing (POCT) has emerged as a critical strategy for rapid diagnostics and monitoring, particularly in low-resource settings.^1-2^ Enhancing the efficiency of POCT for the detection of microorganisms necessitates the integration of modern advancements in DNA and RNA recognition, as well as isothermal amplification techniques.^3-5^ Of particular relevance for practical applications is the utilization of a multifaceted approach that combines high sensitivity and specificity in molecular genetic recognition of nucleic acids at a constant temperature with straightforward and expedient detection of the resultant products. Prominent molecular genetic tools include CRISPR (Clustered Regularly Interspaced Short Palindromic Repeats)/Cas (CRISPR-associated protein). CRISPR/Cas-based diagnostics, leveraging guide RNA (gRNA)-directed nucleases, have emerged as powerful tools for nucleic acid detection due to their exceptional specificity and intrinsic signal amplification via nonspecific trans-nuclease (collateral) activity of Cas proteins.^6-8^ DNA/RNA target recognition is determined by the formation of a ternary complex of Cas–gRNA–target, requiring a sequence of nucleic acid target complementary to the gRNA. Upon successful target recognition, Cas cleaves the target and acquires additional indiscriminate trans-nuclease activity. Specifically, enzymes such as Cas12a and Cas14a demonstrate trans-nuclease activity towards single-stranded (ss) DNA, whereas Cas13a is specific for ssRNA substrates.^9^ In diagnostic systems utilizing Cas proteins with trans-nuclease activity, the cleaved oligonucleotide is modified with labels (referred to as a probe), with the detected signal derived from the cleaved or uncleaved state of the probe. The choice of labels is determined by the used detection method, among which the received the main recognition are instrumental fluorescence detection based on a fluorescent label/quencher pair and instrumental-free detection using paper-based methods.^6, 10^

Lateral flow tests (LFTs) are the primary tool for instrument-free detection in CRISPR/Cas-based diagnostics among paper-based methods^11-12^ largely due to their ease of use. The detection process for LFTs begins with immersing the test strip in a liquid sample. Capillary forces then draw the liquid through the strip, passing through the zones with the labeled conjugate, the test zone, and the control zone, which contains immobilized biomolecules. The detected signal is generated by the capture of the conjugate with a detectable label in both the test and control zones. Specifically, the signal in the test zone depends on the presence of the analyte. For instance, in the sandwich format of an LFT, a signal appears in the test zone only when a complex is formed between a biomolecule in the test zone, the analyte, and a conjugate with a label.^13-14^ In the case of the Cas with trans-nuclease activity, the analyte detected by the LFT is a probe. The most widely adopted CRISPR/Cas–LFT strategy is based on the DETECTR (DNA Endonuclease Targeted CRISPR Trans Reporter) format, utilizing an ssDNA probe labeled with fluorescein (FAM) and biotin (Bio) at opposite ends.^15^ Both FAM and Bio are low-molecular-weight ligands that have well-established high-affinity and specific receptors: streptavidin for Bio, and antibodies for FAM (antiFAM). However, in conventional LFT for detection of FAM-ssDNA-Bio, test and control zones have an atypical arrangement. The first zone in the direction of fluid flow is referred to as the control zone, where streptavidin is adsorbed. The second zone is the test zone, which contains antibodies specific to immunoglobulins (or protein A). These antibodies bind to the conjugate of gold nanoparticles (GNPs) and antiFAM, as depicted in **Figure 1A**. Consequently, when FAM-ssDNA-Bio remains intact and is not cleaved by Cas12a, it forms a colored ternary complex (streptavidin – FAM-ssDNA-Bio – antiFAM-GNP) in the control zone. Therefore, antiFAM-GNP conjugate does not migrate into the test zone. When Cas12a is activated and cleaves ssDNA, a reduced quantity of the ternary complex is generated in the first control zone. Meanwhile, some of the conjugate, together with the cleaved probe, proceeds to the second test zone, where it is captured and visually detected.

**Figure 1.**
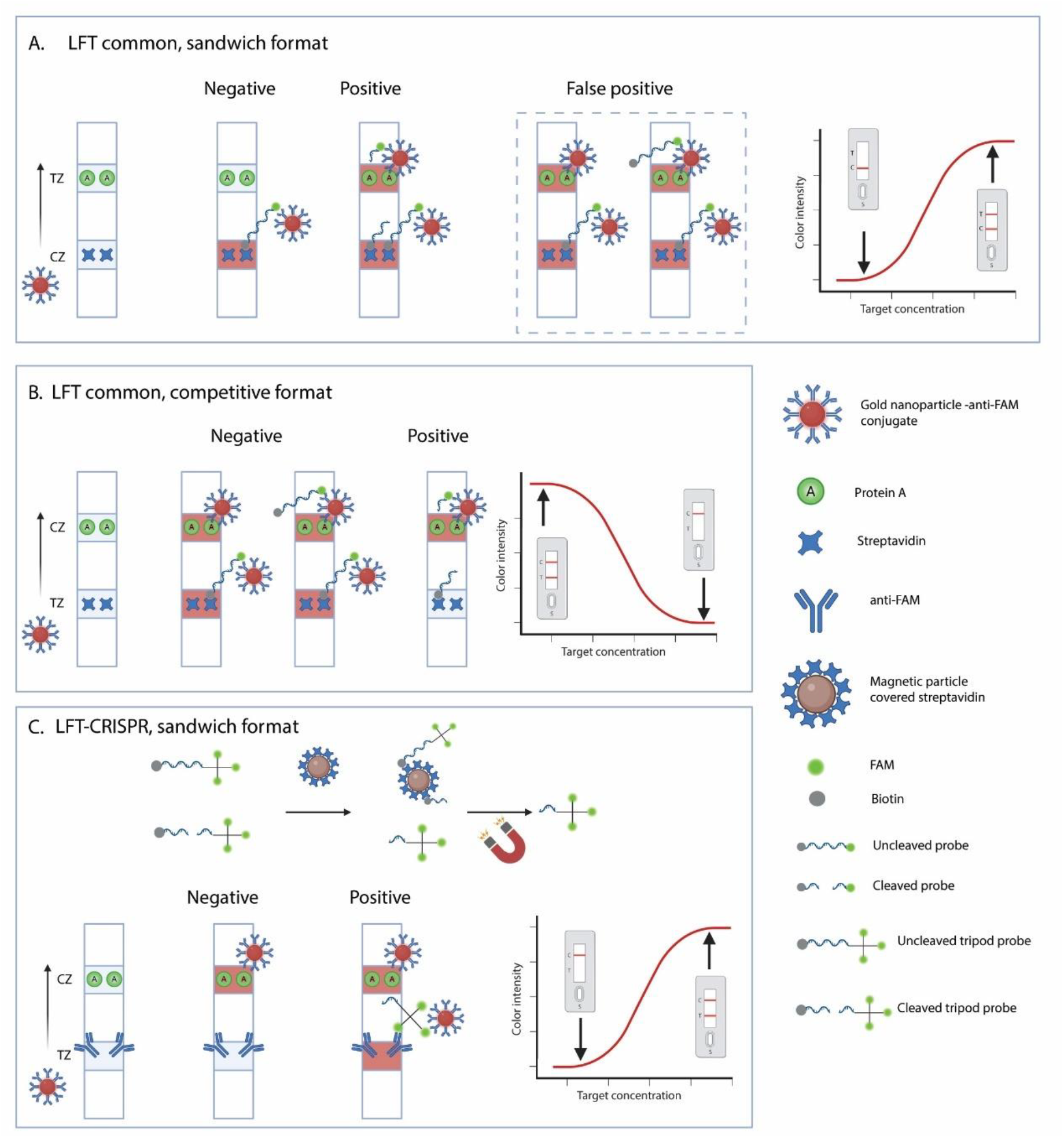
Schemes of test strips for detection of ssDNA probes for CRISPR/Cas: LFT-common **(A), (B)**, LFT-CRISPR **(C)**. TZ is an abbreviation for test zone, CZ is for control zone, FAM is for fluorescein label, and Bio is for biotin. Here and below the schemes were created in BioRender.

Despite the considerable number of biosensors implemented according to the DETECTR scheme,^16-21^ the scheme relies on the indirect dependence of the coloration in the test zone on the cleaved probe. This is due to the fact that the probe itself is not an indispensable component of the detected complex in the test zone. Coloration in the test zone occurs when protein A – antiFAM-GNP conjugate, protein A– antiFAM-GNP conjugate – uncleaved probe, and protein A– antiFAM-GNP conjugate – cleaved probe complexes are formed (the schemes are shown in **Figure 1A**). Therefore, this scheme introduces a significant risk of false-positive results. The capturing of the whole conjugate in the first zone (negative test) may not occur, the conjugate will to migrate to the second zone, where binding can occur independently of the cleaved probe, thus leading to a false-positive result. The reasons for false-positive results can be associated with an initially incorrect balance of reaction components or a change in this balance due to storage of the LFT.

To overcome this design limitation and create alternative combinations of CRISPR/Cas with LFT, several solutions have been proposed. One of the simplest options considers the LFT illustrated in **Figure 1B** as a competitive format.^22-28^ In this format, the binding zone with immobilized streptavidin, which meets the capillary flow first, becomes a test zone, and the second zone for capture of GNP-antiFAM becomes a control zone. The results are interpreted in alignment with the competitive assay format: one colored line (control zone) is a positive result, and two colored-lines correspond to a negative result. In this case, the probe directly affects the coloration in the test zone; however, the dependence of the signal on the dsDNA-target concentration is inverse. In addition, there are schemes that preserve both the direct location of the binding zones (the first is a test zone, the second is a control zone) and the direct dependence of the coloration on target concentration. For instance, DIRECT^2^ (DNA-Immunoglobulin Reporter Endonuclease Cleavage Test), employs a probe that consists of a protein part (immunoglobulin G) and a DNA part (ssDNA) attached to a carrier. When activated, Cas12 cleaves the ssDNA, releasing the protein component, which can then be detected by LFT.^29^ Another example is CLIPON (CRISPR and Large DNA assembly Induced Pregnancy strips for signal-ON detection), which also utilizes an ssDNA-protein conjugate as a probe. In this case, the protein – human chorionic gonadotropin is detected by pregnancy test strip if Cas12a cleavages the ssDNA.^30^ Also, a ready-made pregnancy test and a conjugate of GNPs with poly(A) ssDNA and goat antimouse IgG were used for BE-CATCH strategy that provided sensitivity, specificity and user-friendliness.^31^ However, such hybrid probes pose challenges in terms of synthesis and stability. Therefore, the aim of this research was to develop a convenient universal oligonucleotide probe that can be easily synthesized in large amounts. This probe should reliably produce a color change in the test zone, be part of the ternary complex in the test zone only when cleavage occurs, and provide a direct relationship between color intensity and the concentration of dsDNA-targets.

Here, we developed a novel type of DNA probe (polyFAM probe) which feature multiple FAM labels and an ssDNA labeled with Bio. This design combines all the essential benefits for CRISPR/Cas12a – LFT. As a result of comparing five different polyFAM probes, we selected a highly effective tripod probe that contains three FAM labels. Upon Cas12a-mediated cleavage, a three-label fragment is released, enabling detection via a direct sandwich LFT with antiFAM antibodies in both the test zone and gold nanoparticle conjugate (LFT-CRISPR). To ensure specificity of the signal, uncleaved probes are removed via magnetic separation, utilizing a Bio-label and streptavidin-coated magnetic particles (MP-Str). The principle of this tripod-probe – LFT strategy is illustrated in **Figure 1C**. A priority foodborne pathogen, *Salmonella* Typhimurium, which poses a significant health risk worldwide,^32-33^ was chosen for testing the probes. For *S*. Typhimurium, it is important to minimize analysis time for rapid detection, especially after a brief enrichment phase to grow the pathogen. Additionally, higher sensitivity in the analysis reduces the time needed for pathogen growth.^34^

## EXPERIMENTAL SECTION

### Materials and Reagents

All of the oligonucleotide sequences (**Tables S1**), chemicals and materials, preparation of DNA targets, and loop-mediated isothermal amplification are described in **Section 1, Supporting Information**.

### Preparation of lateral flow test strips for probe detection

Gold nanoparticles (GNPs) were synthesized via sodium citrate reduction of Au^4+^, according to the Frens method.^35^ Conjugate of synthesized GNPs and antiFAM antibodies was prepared using physical adsorption as described by Safenkova et al.^36^ Details on the synthesis and characterization of GNPs and conjugates via spectrophotometry, transmission electron microscopy (TEM), and dynamic light scattering (DLS) are provided in **Section 2, Supporting Information**.

Two types of test strips were made (see **Figure 1b,c**), differing in the test zone component and the applied conjugate: (1) streptavidin immobilized in the test zone with GNP-antiFAM conjugate (LFT-common) and (2) anti-FAM immobilized in the test zone with GNP-antiFAM conjugate (LFT-CRISPR). The control zone was formed using protein A. Reagents for the test and control zones were dispensed onto CN-95 nitrocellulose membranes at 0.15 µL/mm (1 mg/mL of anti-FAM, 1.0 mg/mL of streptavidin, 0.5 mg/mL protein A) using an IsoFlow multichannel dispenser (Imagene Technology, St. Lebanon, NH, USA). Conjugates were applied to PT-R5 glass fiber membranes at 1.6 µL/mm, pre-adjusted to an optical density of A_520_ = 3.0. The nitrocellulose and glass fiber membranes were dried for 2 hours at 37 °C, then assembled into a multimembrane composite on a plastic backing, consisting of a nitrocellulose membrane, conjugate pad, sample pad, and absorbent pad. The membrane composite was cut into 3 mm-wide strips using an automatic guillotine ZQ2002 (Shanghai Kinbio Tech, Shanghai, China), sealed in airtight packaging with silica gel, and stored at room temperature.

### Testing with lateral flow test strips

Test strips were vertically immersed in a sample of at least 100 µL for 10 minutes. The results were visually detected and digitally recorded by scanning (Canon 9000F Mark II scanner, Canon, Tokyo, Japan) and processing the densitometric signal using TotalLab Quant software (TotalLab Great Britain, Newcastle upon Tyne, UK). Each sample was analyzed in at least two – three replicates, calculating the mean value and standard deviation (SD). The visual limit of detection (vLOD) was determined as the minimal target concentration at which the color intensity in the test zone was visible to the naked eye that corresponded to 1 relative unit (RU) for the used procedures of image processing (Panferov et al. 2018). The digital detection limit was determined using the 3σ method.

### Preparation of polyFAM probes consists of dsDNA with multiple FAM labels and an ssDNA labeled Bio

PolyFAM DNA probes were generated by incorporating FAM-11-dCTP during PCR amplification. The DNA template for the 160 bp amplicon was enhanced green fluorescent protein (eGFP) gene fragment within the pEGFPN1 plasmid (**Table S1, Supporting Information**). The reaction mixture included Taq buffer, 60 µM dNTPs, 500 nM forward primer (eGFP-PCR-F) with a 15dT tail and a 5’-biotin label, 500 nM reverse primer (eGFP-PCR-R) with a 5’-FAM label (primers are presented **Table S1, Section 1, Supporting Information**), 0.75 ng/µL pEGFPN1, and 0.1 U/µL Taq DNA polymerase, following the method described in (Ivanov et al. 2023). To obtain a DNA probe with polyFAM and a single biotin label, FAM-11-dCTP was added at different concentrations: 10, 20, 40, and 60 µM. Thus, dCTP was proportionally reduced, keeping the total amount of FAM-11-dCTP and dCTP equal to 60 µM. PCR was performed for 35 cycles (denaturation at 95 °C for 30 s, annealing at 65 °C for 30 s, and elongation at 72 °C for 1 min) in a T100 thermal cycler (Bio-Rad, Hercules, CA, USA). The PCR product was concentrated using Amicon Ultra 3K (Merck Millipore, Burlington, MA, USA) and separated by electrophoresis in a 2% agarose gel in 20 mM Tris-acetate buffer with 0.2 mM EDTA, pH 8.3 (TAE) by electrophoresis (horizontal chamber from Helikon, Russia; power source from Bio-Rad, USA), stained with ethidium bromide. DNA bands were visualized under UV light using a GenoSens 2150 GelDoc system (Clinx Science Instruments, Shanghai, China). The polyFAM DNA amplicons were cut out from agarose gel and purified using the Cleanup Standard kit. DNA concentration was measured using a Nanodrop ND-2000 spectrophotometer (ThermoFisher Scientific, Waltham, MA, USA).

### CRISPR/Cas12a reaction with fluorescent detection

For the CRISPR/Cas12a reaction, a mixture containing 60 nM gRNA (see **Table S1, Section 1, Supporting Information**) and 67 nM Cas12a (NEB, USA) in a buffer consisting of 10 mM Tris-HCl (pH 7.9), 50 mM NaCl, 10 mM MgCl_2_, and 100 µg/mL BSA (NEBuffer r2.1, New England Biolabs, USA) was prepared and incubated at 25 °C for 10 min. Subsequently, the ROX-dT15-BHQ2 probe was added to a final concentration of 500 nM, followed by the addition of 2 µL of the sample (DNA amplicon, or LAMP reaction mixture without SYBR Green I), bringing the total reaction volume to 30 µL. Fluorescence detection of the ROX-dT15-BHQ2 probe was performed kinetically using a Light Cycler 96 system (Roche, Switzerland) over 40 min with λ_ex_ = 578 nm and λ_em_ = 604 nm.

### CRISPR/Cas12a reaction using polyFAM probe and LFT detection

The CRISPR/Cas12a reaction was performed as described in **Section “CRISPR/Cas12a reaction with fluorescent detection”**, except that a polyFAM probe was added to a final concentration of 33 nM instead of the ROX-dT15-BHQ2 probe, maintaining a total reaction volume of 30 µL. After 30 min of reaction time, the mixture was incubated with MP-St for 5 min at 25 °C. Magnetic separation was used to collect MP-St-bound biotinylated components, and the supernatant (30 µL) was transferred to a tube containing 30 µL of PBSTx2. The CRISPR-LFT strip was immersed in the solution, and the results were visually assessed after 10 min. For digital analysis, test strips were scanned, and signal quantification was performed as described in **Section “Testing with lateral flow test strips”**.

## RESULT AND DISCUSSION

### Concept of polyFAM probes and their preparation

The main idea behind the proposed polyFAM probes is to utilize a structure that incorporates multiple FAM labels along with a single biotin-labeled ssDNA fragment, which undergoes trans-nuclease cleavage upon activation by Cas12a. The Bio label is essential for the separation of the uncleaved polyFAM probe using MP-Str and magnet (see **Figure 1C**). Effective separation is crucial for the success of this approach, which is why we selected the Bio-streptavidin pair, known for its exceptionally high binding constant (up to 10^−15^ M).^37^

The number of FAM labels in the polyFAM should be sufficient to ensure effective formation of a ternary complex in the LFT-CRISPR test zone (antiFAM immobilized in the test zone – polyFAM probe – antiFAM-GNP conjugate), as shown in **Figure 1C**. To address this requirement, we designed two types of polyFAM probes.

The first type of polyFAM probe is the tripod probe, which is a single-stranded dT probe featuring a triple branch point at the 5′-end due to the use of trebler phosphoramidite (the structure is presented in **Figure S3, Section 3, Supporting Information**) during oligonucleotide synthesis. Each branch extends through a hexaethylene glycol (Heg) spacer (approximately 1.7 nm in length) and terminates with a FAM label at the 5′-end. Consequently, each tripod probe contains three FAM labels at the 5′-end and one Bio label at the 3′-end. This probe can be easily synthesized using standard oligonucleotide synthesis techniques with trebler phosphoramidite. We designed two variants of the tripod probe: one with fifteen thymine nucleotides (tripod-dT15) and another with fifty thymine nucleotides (tripod-dT50) (**Figure 2A, Table S1, Section 1, Supporting Information**). As a control, we also used a probe featuring a single FAM label and a single Bio label (FAM-dT15-Bio).

**Figure 2.**
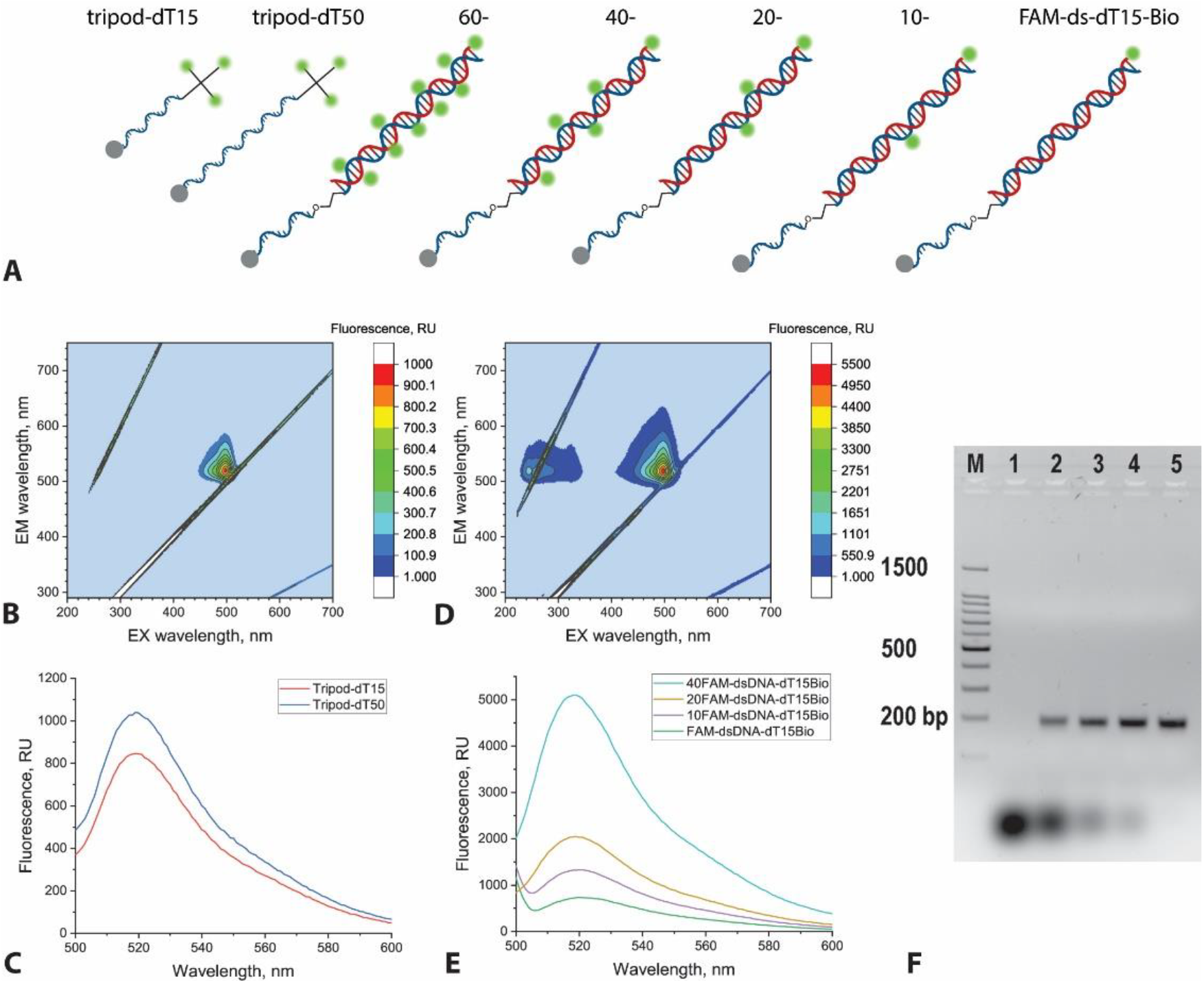
PolyFAM probes and their characteristics: schemes of polyFAM probes **(A)**, excitation-emission matrixes of 50 nM tripod-dT15 **(B)** and 50 nM 40FAM-dsDNA-dT15Bio **(D)**, fluorescence spectra at λex = 497 nm for tripod probes **(C)** and polyFAM-dsDNA-T15Bio probes **(E)**, electrophoresis of the obtained polyFAM probes (60FAM-dsDNA-dT15Bio – 1, 40FAM-dsDNA-dT15Bio – 2, 20FAM-dsDNA-dT15Bio – 3, 10FAM-dsDNA-dT15Bio – 4, FAM-dsDNA-dT15Bio – 5) **(F)**.

The second type of polyFAM probe is the polyFAM-dsDNA-dT15-Bio probe (**Figure 2A**). This is a double-stranded DNA fragment produced through PCR, which incorporates multiple FAM labels due to the use of a labeled cytosine derivative (FAM-11-dCTP). We selected a fragment length of 160 bp because previous studies have demonstrated that this length allows for effective binding with test strips, even with only a pair of labels.^36^ The nucleotide sequence of the dsDNA fragment corresponds to a portion of the eGFP gene (see **Table S1, Section 1, Supporting Information**). To modulate the number of FAM labels in the synthesized probe, we varied the concentration ratio of dCTP (µM) / FAM-11-dCTP (µM) during PCR: 60/0, 50/10, 40/20, 20/40, 0/60. The Bio label was introduced via a primer containing a C3 spacer between the annealing sequence and dT15-Bio. The C3 spacer causes the polymerase to stop, resulting in a dT15-Bio single-stranded overhang on the double-stranded amplicon. This method produced a dsDNA construct with polyFAM labels and an ssDNA dT15-Bio tail. Electrophoretic analysis of the PCR amplicons revealed a single product close to 200 bp (**Figure 2F**). The visible increase in size beyond the 160 bp of eGFP-160 template (**Table S1, Supporting Information**) probably occurs due to the inclusion of a single-strand 15dT and labels in the amplicon. Moreover, when dCTP is completely replaced by FAM-11-dCTP, the amplicon is almost not formed. The yield of the PCR product improves as the ratio of dCTP to FAM-11-dCTP increases (**Figure 2F**).

A comparison of fluorescence intensities among the obtained probes confirmed varying degrees of FAM incorporation: three per tripod probes and two, three, and nine FAMs per polyFAM-dsDNA-dT15-Bio probes (**Figure 2B-E**).

### Testing of polyFAM probes using lateral flow test strips

To evaluate the performance of the proposed probes, two types of test strips were prepared (see **Figure 1**):

1. **LFT-common**: This test strip is based on the conventional design, with streptavidin immobilized in the test zone and an antiFAM-GNP conjugate. The LFT-common configuration was used to characterize probe binding via FAM and Bio-labels. It is particularly useful for assessing the efficiency of probe cleavage, which results in the cutting off of the Bio label, thereby preventing detection with the LFT-common.
2. **LFT-CRISPR**: Proposed in this study, this test strip has antiFAM immobilized in the test zone and an antiFAM-GNP conjugate. This configuration allows for the detection of the polyFAM part of the probe in a sandwich assay format.

The characteristics of the synthetized GNP and antiFAM-GNP conjugates are detailed in **Section 2** of **Supporting Information**.

Initially, the probes were compared due to their binding in the test zone via FAM and Bio labels using the LFT-common. The concentration-dependent responses for single-label probes (FAM-dT15-Bio and FAM-dsDNA-dT15-Bio) and polyFAM-labeled probes (tripod-dT15, tripod-dT50; 10-, 20-, and 40FAM-dsDNA-dT15-Bio) in test zones of LFT-common revealed that the vLOD for all probes was 0.1-0.2 nM (**Figure S4, Section 4, Supporting Information**). These findings confirm that all proposed probes effectively bind to LFT-common via both FAM and Bio-labels. Moreover, LFT-common is an effective tool to assess probe cleavage in the CRISPR/Cas12a reaction because cleavage of the dT-strand carrying the Bio-label reduces binding in the test zone.

For probe detection using LFT-CRISPR, several FAM labels are required. Testing the polyFAM probe indicated binding for all probe types (**Figure S5, Section 4, Supporting Information**). Therefore, even tripod probe with three FAM labels at one end of the probe—spaced up to 3.5 nm apart, on calculations of linker length for two Heg spacers—are adequate for forming of a ternary complex in the test zone (antiFAM – probe –antiFAM-GNP conjugate). At high tripod probe concentrations (50 nM, equivalent to 5 pmol per reaction), a pronounced hook effect was observed, characterized by a significant decrease in test zone intensity (see **Figure S5**). The hook effect likely arises due to probe oversaturation of both the test zone (3 pmol antiFAM) and the conjugate surface (approximately 0.016 pmol conjugate per test strip), which interferes with ternary complex formation in the test zone. The concentrations at which the hook effect occurs determine the upper limit of the concentration range for the polyFAM probes in the CRISPR/Cas12a reaction. Differences in probe detection limits were more apparent, with the vLOD decreasing in the following order: 10FAM-dsDNA-dT15-Bio = 20FAM-dsDNA-dT15-Bio (1 nM) > tripod-dT15 (0.6 nM) > 40FAM-dsDNA-dT15-Bio (0.4 nM) > tripod-dT50 (0.2 nM). These results enabled the implementation of all polyFAM probes following cleavage in the CRISPR/Cas12a reaction for detection using LFT-CRISPR.

### Testing of polyFAM probes in the CRISPR/Cas12 reaction

To implement the CRISPR/Cas12 assay with a polyFAM probe and LFT-CRISPR, two primary requirements must be realized. First, following the CRISPR/Cas12a reaction, the concentration of the cleaved probe should fall within the working range, i.e., 0.6–17 nM for tripod-dT15, 0.2–17 nM for tripod-dT50, and 1–10 nM for polyFAM-dsDNA-dT15-Bio. Second, after magnetic separation of the uncleaved probe, LFT-CRISPR testing should show no signal in the test zone.

For the comparison of probes in the CRISPR/Cas12a reaction, we selected the invA gene of *S*. Typhimurium as the target. This gene is a commonly used specific marker in PCR-based and isothermal amplification-based diagnostic methods.^34, 38-39^ A 959 bp dsDNA fragment, obtained through PCR and purified (sequence is presented in **Table S1**, electrophoresis data is provided in **Figure S6, Supporting Information**), was used as the dsDNA-target recognized by the gRNA, which activates the trans-nuclease activity of Cas12a.

For each probe, we performed the CRISPR/Cas12a reaction using 0.5 nM and 0 nM dsDNA-target, along with 33 nM probe. This setup ensures a final probe concentration of 10 nM in the solution where the test strip is immersed. Under these conditions, we initially tested the samples with LFT-common to evaluate cleavage efficiency for different probes. The results indicate that at zero dsDNA*-* target concentration, the test zones demonstrate strong coloration (**Figure 3A,C**). However, at 0.5 nM dsDNA-target, there was no coloration only for tripod-dT50 (approximately 100% cleavage), and a strong decrease in coloration was observed for tripod-dT15, indicating ∼90% cleavage of tripod-dT15 was cleaved. In contrast, the polyFAM-dsDNA-dT15Bio probes showed minimal decrease in coloration, around 10%, suggesting that approximately 20-30% cleavage of these probes (according to the concentration dependences in **Figure S4**). These results indicate that the cleavage efficiency of tripod probes is significantly higher than that of polyFAM-dsDNA-dT15Bio probes.

**Figure 3.**
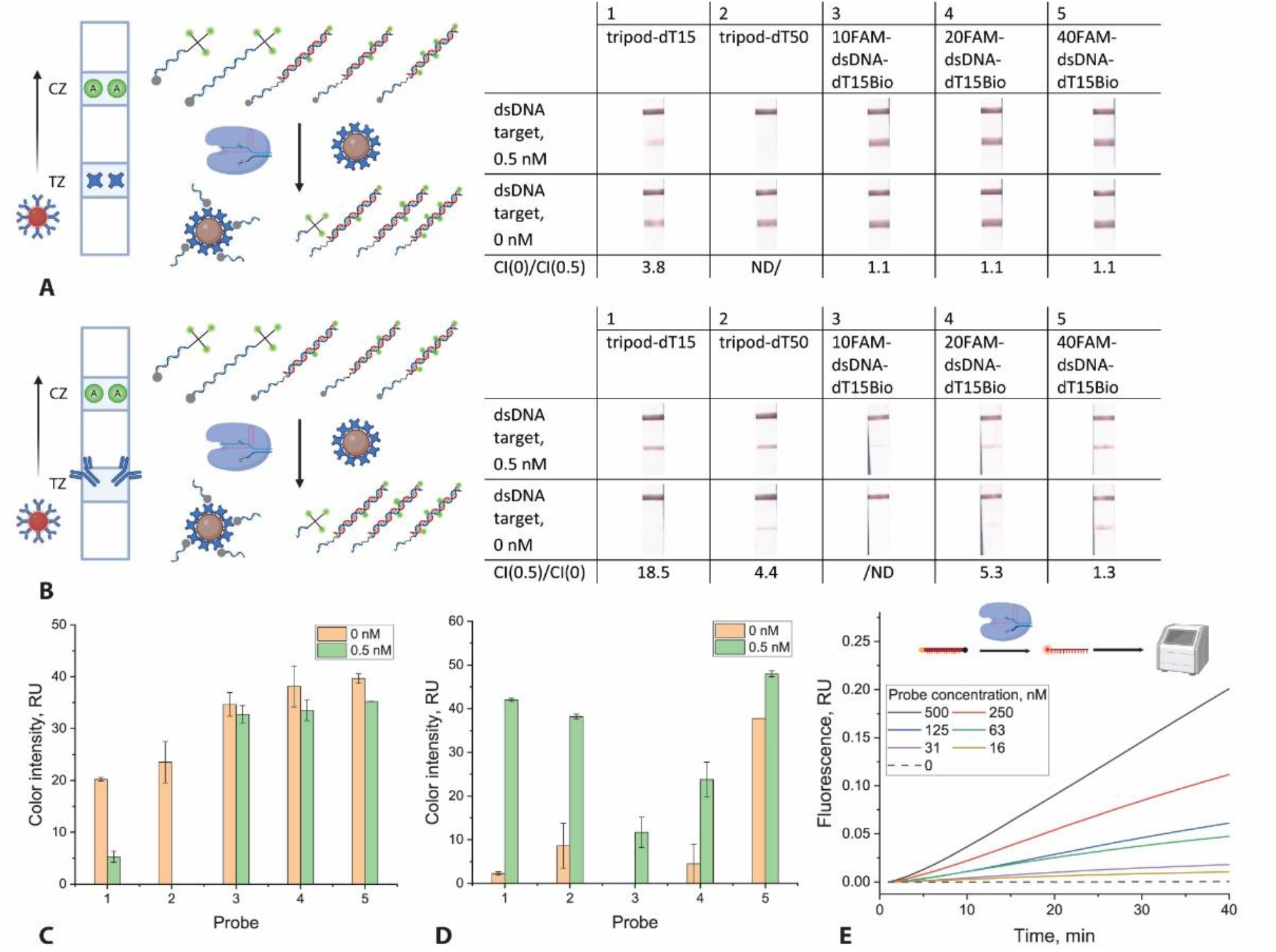
Results of CRISPR-Cas12a reactions for 0.5 nM / 0 nM dsDNA-target with different polyFAM probes and LFT-common **(A), (C)** or LFT-CRISPR **(B), (D)** as well as with ROX-dT15-BHQ2 at different concentrations (from 500 to 16 nM) with fluorescence detection **(E)**. ND means that the signal was not detected.

Second, magnetic separation was performed for 5 minutes to remove uncleaved probes, after which the samples were analyzed using LFT-CRISPR. Test zone coloration was observed for all probes at a concentration of 0.5 nM dsDNA-target (**Figure 3B,D**). However, for 10- and 20FAM-dsDNA-dT15Bio probes, the coloration in the test zone was relatively weak, likely due to low cleavage efficiency (see **Figure 3A,C**). An important factor was the absence of visually detectable coloration (< 1 RU) in the test zone at zero dsDNA-target concentration, which was characteristic of the tripod-dT15 and 10FAM-dsDNA-dT15Bio probes (indicated by orange bars in **Figure 3D**). Therefore, among all proposed probes, tripod-dT15 fully meets the required criteria for Cas12a-based analysis with LFT-CRISPR detection. Additional tests, including evaluations with various combinations of CRISPR/Cas12a reaction components, confirmed the suitability of the tripod-dT15 probe (results are provided in **Table S2, Section 6, Supporting Information**). Moreover, tripod-dT15 is advantageous due to its simpler synthesis, as it can be obtained via commercial synthesis.

To assess the efficiency of the polyFAM probes and the performance of LFT-CRISPR, we used a standard probe that features a fluorophore and quencher at opposite ends (ROX-dT15-BHQ2, see **Table S1, Supporting Information**). Fluorescence detection was employed to measure the signal from the cleaved probes. These systems typically require probes at considerably higher concentrations (around 500 nM). We performed a reaction with 0.5 nM dsDNA*-*target and tested various concentrations of ROX-dT15-BHQ2 (500, 250, 125, 63, 31, and 16 nM), which revealed significant differences in fluorescence signal intensity (**Figure 3E**). Consequently, we found that the optimal probe concentration for fluorescence-based detection (ROX-dT15-BHQ2) is approximately an order of magnitude greater than that for LFT-CRISPR (tripod-dT15).

### Analytical capabilities of the CRISPR/Cas12a system with the tripod-dT15 probe and LFT-CRISPR

The CRISPR/Cas12a test systems show a nearly linear increase in reaction products over time as well illustrated in **Figure 3E**. A single activated Cas12a can cleave up to 17 oligonucleotides per second, depending on the nuclease type and the characteristics of the oligonucleotides.^40^ This turnover rate often necessitates the use of pre-amplification methods, such as loop-mediated isothermal amplification (LAMP) or other isothermal amplification techniques, to enhance the sensitivity of the assays.^41^ However, several amplification-free CRISPR/Cas12a strategies and test systems based on them have been developed,^42^ indicating that the tripod-dT15 probe and LFT-CRISPR could play a significant role in improving amplification-free approaches. Considering these factors, we evaluated the analytical performance of the CRISPR/Cas12a system using the tripod-dT15 probe and LFT-CRISPR in two scenarios: direct detection of dsDNA targets without prior amplification and detection of dsDNA targets following LAMP.

For direct target DNA detection without prior amplification, we used the dsDNA-target (invA gene, 959 bp) at concentrations ranging from 1 nM to 0.25 pM. The resulting vLOD was determined to be 1.4 pM (2 × 10^7^ copies/reaction) for dsDNA-target (**Figure 4A,B**). This value is 500-fold lower than previously reported for another test system, termed DIRECT^2^, which utilizes an IgG-DNA probe and a direct concentration-dependent signal on the test strip.^29^ The significant decrease in vLOD may be attributed to two main factors: a more efficient gRNA – dsDNA-target pair and/or a more effective probe. To evaluate the impact of the gRNA – dsDNA-target interaction, we repeated the CRISPR/Cas12a reaction using conventional fluorescence detection with the ROX-dT15-BHQ2 probe. The detection limit in this case was found to be 10 pM, which is still lower than the limit of detection reported for fluorophore/quencher-based detection,^29^ but only by a factor of 50. Hence, both factors contribute to the high sensitivity of tripod-dT15–LFT-CRISPR for equipment-free visual detection of CRISPR/Cas12a reactions.

**Figure 4.**
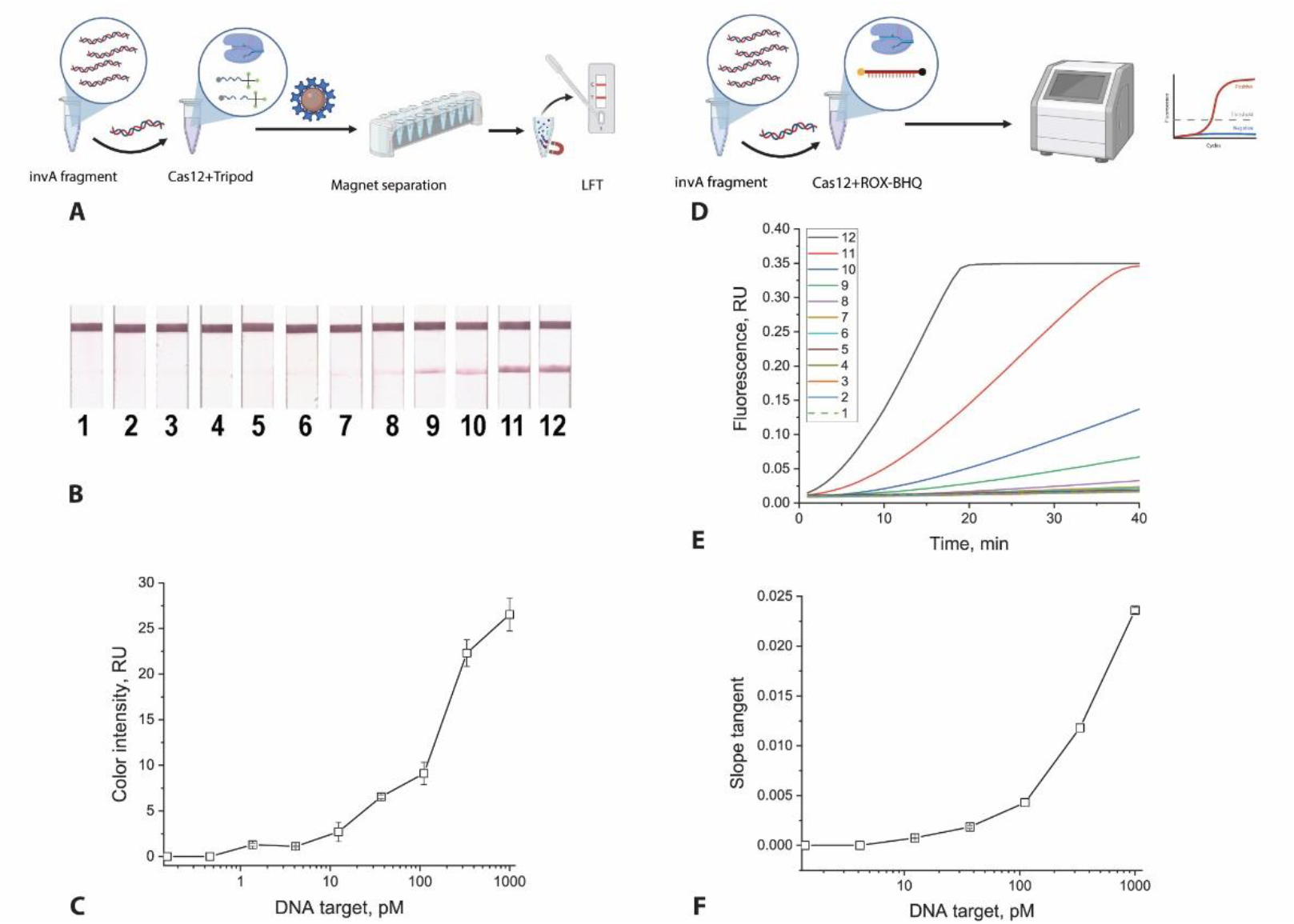
CRISPR/Cas12a without preamplification for dsDNA (959 bp of the invA gene of *S*. Typhimurium), where numbers indicate: 1 – 0, 2 – 0.02, 3 – 0.05 ng/µL, 4 – 0.2, 5 – 0.5, 6 – 1.4, 7 – 4, 8 – 12, 9 – 37, 10 – 111, 11 – 333, 12 – 1000 pM. Scheme of Cas12a–tripod-dT15–LFT-CRISPR **(A)**, test strips for tripod-dT15 – LFT-CRISPR **(B)**, concentration dependence based on color intensities of test zones **(C)**. Scheme of CRISPR/Cas12a with ROX-dT15-BHQ2 and fluorescence detection **(D)**, fluorescence curves obtained for ROX-dT15-BHQ2 **(E)**, concentration dependence based on fluorescence intensity at 40 minutes **(F)**.

For the first time, the obtained result demonstrates that equipment-free visual detection using LFT can achieve approximately 1 pM and be an order-of-magnitude lower detection limit compared to fluorophore/quencher-based detection. This result holds promise for the development of amplification-free CRISPR/Cas12a test systems for DNA targets that require this level of sensitivity.

### CRISPR/Cas12a with tripod-dT15 probe and LFT-CRISPR combined with LAMP

Detection of dsDNA-targets in a CRISPR/Cas12a reaction following LAMP involves the identification of LAMP amplicons, which are concatemeric structures containing repeated fragments of the selected DNA targets (gel electrophoresis scans confirming the heterogeneity of amplicons, fluorescence curves of SYBR Green I staining for target and non-target cells are presented in **Figure S7, Supporting Information**). Therefore, if the DNA target initiates the LAMP reaction, the resulting amplicons are present in significantly higher amounts than the limit of detection (LOD) for CRISPR/Cas12a. A comparative evaluation of LAMP products obtained from different target concentrations (ranging from 2 × 10^9^ to 3 copies per reaction) demonstrated that the reliably detectable product using SYBR Green staining was 740 copies/reaction (fluorescence curves of LAMP and their concentration dependence are provided in **Figure S8, Supporting Information**).

Integrating LAMP with a subsequent CRISPR/Cas12a reaction enhances both sensitivity and specificity.^6, 10^ Therefore, all mixtures of LAMP reactions were used as samples for the CRISPR/Cas12a reaction. The testing was performed in two formats: fluorescence detection using ROX-dT15-BHQ2 and lateral flow detection using tripod-dT15–LFT-CRISPR. Both formats provided increasing analytical sensitivity compared to LAMP alone, with a detection limit of 27 copies/reaction, which is 27 times lower than LAMP without CRISPR/Cas12a (**Figure 5B**, fluorescence curves for ROX-dT15-BHQ2 are presented in **Figure S9B, Supporting Information**). Several key observations can be made from these results. First, both fluorescence signals and test zone color intensities were strong and uniform across all concentrations, unlike the amplification-free CRISPR/Cas12a setup, where signal intensity varied with concentration (see **Figure 4**). This can be attributed to the nature of LAMP, which generates a large number of DNA amplicons regardless of the initial DNA target concentration. Second, it is likely that LAMP amplification occurs even at concentrations between 27 and 740 copies/reaction, though this was not evident when using SYBR Green staining (see **Figure S8, Supporting Information**). This suggests that at these concentrations, 40 minutes of LAMP produces an insufficient number of amplicon copies for SYBR Green visualization but a sufficient amount of DNA target (>1.4 pM = 2 × 10^7^ copies/reaction) to activate Cas12a and initiate probe cleavage. Third, when preceded by amplification, Cas12a–tripod-dT15–LFT-CRISPR demonstrated analytical sensitivity comparable to Cas12a–ROX-dT15-BHQ2– fluorescence detection. However, unlike the fluorescence-based method, the LFT-based detection does not require specialized equipment and can be implemented in minimally equipped laboratories.

**Figure 5.**
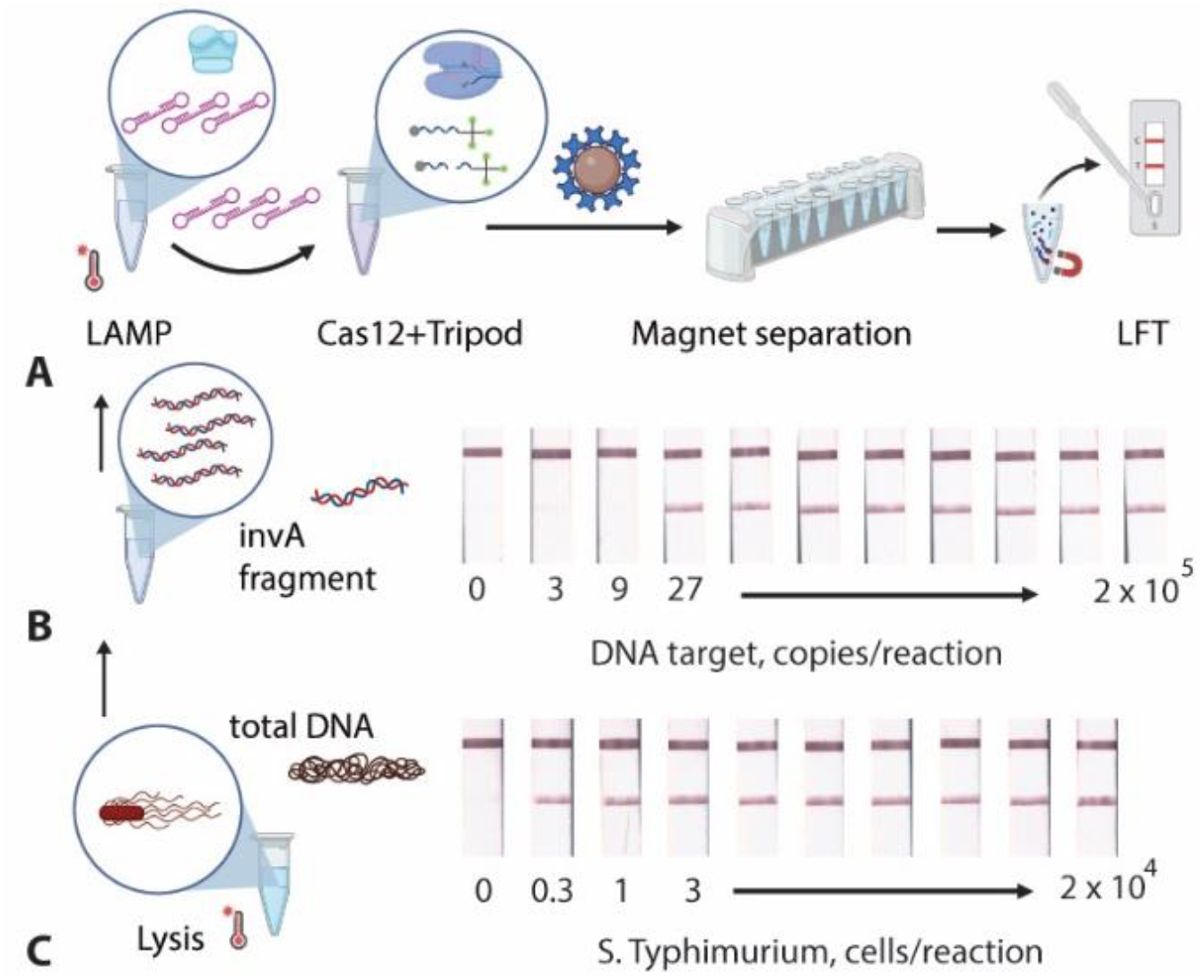
CRISPR/Cas12a with LAMP samples. Scheme of Cas12a–tripod-dT15–LFT-CRISPR **(A)**. Test strips for tripod-dT15 – LFT-CRISPR with LAMP of dsDNA (959 bp, invA gene of *S*. Typhimurium) **(B)**, with LAMP of *S*. Typhimurium cells after thermolysis **(C)**.

To further validate the performance of Cas12a–tripod-dT15–LFT-CRISPR, we tested inactivated *S*. Typhimurium cells. Samples obtained after thermal lysis of cells (ranging from 2 × 10^5^ to 0.3 cells per reaction) for 10 min without isolation of total DNA were used for LAMP and showed a positive signal for all concentrations (fluorescence curves for LAMP are presented in **Figure S10, Supporting Information**). These LAMP mixtures were then used as samples for the CRISPR/Cas12a reaction. Strong positive signals were observed for all nine positive samples in both fluorescence-based (ROX-dT15-BHQ2) and lateral flow-based (tripod-dT15–LFT-CRISPR) formats (**Figure 5C, Figure S9C, Supporting Information**). The detection limit for both formats was 0.3 cells/reaction (150 cells/mL), making this method suitable for detecting *S*. Typhimurium cells following short-term enrichment in appropriate culture media.^34^ Comparison with previously reported CRISPR/Cas12a test systems combined with LFT-based detection and pre-amplification^43-45^ revealed that the proposed tripod-dT15 – LFT-CRISPR demonstrated the same sensitivity but a more reliable and convenient way of detection (comparative data are summarized in **Table S3, Supporting Information**). The key advantage of the Cas12a–tripod-dT15–LFT-CRISPR approach in combination with LAMP is the simple and clear visual readout using lateral flow test strips, which offers sensitivity comparable to instrument-based fluorescence detection. Thus, the obtained results demonstrate the advantages of tripod-dT15 – LFT-CRISPR for both amplification-free CRISPR/Cas12a systems and combined LAMP–CRISPR/Cas12a test systems.

## CONCLUSION

We proposed the concept of polyFAM probes and a specialized LFT-CRISPR specifically designed for probe capture in a sandwich format for use in a CRISPR/Cas12a-based assay. This concept provides several advantages: (i) signal intensity in the test zone directly correlate with the concentration of the cleaved probe, and thus the dsDNA-target concentration; (ii) the cleaved polyFAM probe is part of the complex in the test zone; (iii) the polyFAM probe with a single-stranded DNA tail is an easily synthesized structure; and (iv) LFT-CRISPR is based on widely available reagents (antiFAM, antiFAM-AuNPs, and protein A).

Among the polyFAM probes tested, the tripod-dT15 demonstrated the highest efficiency. This probe comprises 15 thymine nucleotides with a biotin label at the 3′-end and a 5′-end branch point extending into three arms, each consisting of a hexaethylene glycol linker terminating in a FAM-label. The probe can be readily synthesized using standard oligonucleotide synthesis techniques with trebler phosphoramidite. The amplification-free CRISPR/Cas12a assay, based on tripod-dT15 and LFT-CRISPR, achieved a detection limit of 1.4 pM for the DNA target (dsDNA, 959 bp of the invA gene of *S*. Typhimurium), which is an order-of-magnitude lower detection limit compared to fluorophore/quencher-based detection using equipment. The CRISPR/Cas12a assay combined with LAMP, utilizing tripod-dT15 and LFT-CRISPR, exhibited a detection limit of 27 copies of the DNA target per reaction, which was comparable to instrument-based fluorescence detection. The CRISPR/Cas12a–tripod–LFT assay can be easily adapted to detect other DNA targets by modifying the gRNA sequences. These results highlight the potential of the tripod-dT15 and LFT-CRISPR approach for simple, highly sensitive, and equipment-free detection in CRISPR/Cas12a-based diagnostic systems.

## ASSOCIATED CONTENT

### Supporting Information

Supporting Information file (PDF) contains the following sections: Chemicals and materials; description of methods (preparation of DNA targets, and loop-mediated isothermal amplification, synthesis and characterization of gold nanoparticles and their conjugates); structure of trebler phosphoramidite; characteristics of polyFAM probes with LFT-common and LFT-CRISPR; basic check of the CRISPR/Cas12a test system including tripod-dT15 and LFT-CRISPR; results obtained with loop-mediated isothermal amplification; comparison of CRISPR/Cas12a test-system combined with LFT for *Salmonella* detection.

## AUTHOR INFORMATION

## ACKNOWLEDGMENT

The authors are grateful to Dr. Sergey F. Biketov (State Research Center for Applied Microbiology and Biotechnology, Obolensk, Moscow Region, Russia) for providing bacterial cells.

This work was supported by the Russian Science Foundation (N° 23-46-10011)

