## Supporting Information for "A novel tripod probe and lateral flow test to improve CRISPR/Cas12a assay: benefits of branched probe based on trebler phosphoramidite modification"

**Section 1.** Materials, methods, DNA-targets, primers, and probes

**Chemicals and materials.** Tripod probes, labeled oligonucleotides, primers, and gRNA (the sequences are presented in **Table S1, Section 1, Supporting Information**) were custom-synthesized by Lumiprobe (Moscow, Russia). Nucleotides, Taq polymerase, polymerase buffer, DNA markers for electrophoresis, and the Cleanup Standard DNA extraction kit were obtained from Evrogen (Moscow, Russia). Q5 polymerase, Cas12a, and NEB 2.1r buffer were purchased from New England Biolabs (Ipswich, MA, USA). 6-fluorescein derivative of deoxycytidine triphosphate (FAM-11-dCTP) was acquired from Lumiprobe (Moscow, Russia). The BioMaster LAMP SYBR kit for LAMP with SYBR GREEN I detection and the BioMaster LAMP reagent kit for LAMP detection in agarose gel were obtained from Biolabmix (Novosibirsk, Russia). Agarose was sourced from neoFroxx (Einhausen, Germany), while magnetic nanoparticles coated with streptavidin (MP-Str) were purchased from Chemicell (Berlin, Germany). Amicon Ultra 3K filters were obtained from Merck Millipore (Burlington, MA, USA). The total DNA extraction kit "M-Sorb-Vet-DNA" was supplied by Syntol (Moscow, Russia). Hydrogen tetrachloroaurate (III), bovine serum albumin (BSA), sodium azide, glycerol, and ethidium bromide were purchased from Sigma-Aldrich (St. Louis, MO, USA). Streptavidin and protein A were obtained from Imtek (Moscow, Russia), while monoclonal anti-fluorescein antibodies (anti-FAM) were obtained from Bialexa (Moscow, Russia). Membranes for lateral flow test strips were sourced from Sartorius Stedim Biotech, Germany (nitrocellulose membrane CN-95) and Advanced Microdevices, India (conjugate pad PT-R5, sample pad GFB-R4, absorbent pad AP045, and plastic backing LP25). Analytical-grade pure compounds were used for buffer preparation.

Inactivated *Salmonella enterica* serovar *Typhimurium* cells (ATCC 19585), *S.* Enteritidis, *S.* Paratyphi, *S.* Virchow, *S.* Anatum, and non-target bacterial cells (*Escherichia coli* 0157:H7 ATCC 51658, Yersinia enterocolitica ATCC 51658) were provided by the State Collection of Pathogenic Microorganisms and Cell Cultures “GKPM–OBOLENSK”, State Research Center for Applied Microbiology and Biotechnology (Obolensk, Moscow region, Russia).

**Preparation of DNA targets.** Total DNA was extracted from inactivated *S. Typhimurium* cells using the "M-Sorb-Vet-DNA" kit according to the manufacturer's protocol. DNA concentration was measured using a Nanodrop ND-2000 (ThermoFischer Scientific, USA). The extracted DNA was used as a template to generate a 959 bp PCR amplicon with 959 bp length based on the gene for invasion protein A (invA, GenBank accession NC_003197). For this purpose, 32 PCR cycles were performed in a T100 amplifier (Bio-Rad, USA), each cycle included 30 s denaturation at 95 °C, 30 s annealing at 50 °C, and 3 min elongation at 72 °C. The final reaction mixture contained 0.025 U/µL Q5 polymerase, Q5 buffer, 750 µM dNTP, 500 nM forward primer invA-PCR-F, 500 nM reverse primer invA-PCR-R (**Table S1, Section 1, Supporting Information**), 0.1 ng /µL of total DNA isolated from *S.* Typhimurium. The PCR product was concentrated using Amicon Ultra 3K (Merck Millipore, Burlington, MA, USA) and separated by electrophoresis in a 2% agarose gel in 20 mM Tris-acetate buffer with 0.2 mM EDTA, pH 8.3 (TAE) by electrophoresis (horizontal chamber from Helikon, Russia; power source from Bio-Rad, USA), stained with ethidium bromide. DNA bands were visualized under UV light using a GenoSens 2150 GelDoc system (Clinx Science Instruments, Shanghai, China). The polyFAM DNA amplicons were cut out from agarose gel and purified using the Cleanup Standard kit. DNA concentration was measured using a Nanodrop ND-2000 spectrophotometer (ThermoFisher Scientific, Waltham, MA, USA).

**Loop-Mediated Isothermal Amplification (LAMP).** LAMP amplification was performed using the Biomaster LAMP SYBR kit (Biolabmix, Russia) with SYBR Green I detection and the BioMaster LAMP kit (Biolabmix, Russia) for agarose gel detection (Biolabmix, Russia), according to the manufacturer's instructions. The reaction mixture (25 µL of the final amplification mixture) contained 0.75 U Bst DNA polymerase, 50 mM Tris-HCl (pH 8.9), 10 mM KCl, 1 mM dNTP, 6 mM $MgCl_2$, and 0.25% Tween 20. Primers proposed by (Xia et al. 2021) were added at the following final concentrations: 200 nM invA-LAMP-F3,

200 nM invA-LAMP-B3, 1.6 µM invA-LAMP-FIP, 1.6 µM invA-LAMP-BIP, 400 nM invA-LAMP-LF, and 400 nM invA-LAMP-LB (see sequences in **Table S1, Section 1, Supporting Information**). The 959 bp DNA amplicon or *S. Typhimurium* cells after thermal lysis for 10 minutes at 95 °C were used as DNA templates (2 µL per reaction). Negative controls included water and total DNA from non-target bacterial cells. The reaction mixture was incubated at 65 °C for 40 min.

For real-time fluorescence detection with the Biomaster LAMP SYBR kit, a Light Cycler 96 system (Roche, Rotkreuz, Switzerland) was used with fluorescence detection every minute at the excitation wavelength ($\lambda_{ex}$) of 497 nm and the excitation wavelength ($\lambda_{em}$) of 520 nm. For the BioMaster LAMP kit, incubation was performed using a TDB-120 Dry Block Thermostat (Biosan, Riga, Latvia). The LAMP products were analyzed by horizontal electrophoresis in a 2% agarose gel in TAE buffer stained with ethidium bromide. Gel was documented and analyzed using UV illumination using the GenoSens 2150 GelDoc system (Clinx Science Instruments, China).

**Table S1.** Sequences of the DNA-targets, primers, and probes used in this research

| # | Name | Sequence 5′-3′ | Purpose |
|---|---|---|---|
| 1 | eGFP-160 | CGCTACCCCGACCACATGAAGCAGCACGACTTCTTCAAGTCC GCCATGCCCGAAGGCTACGTCCAGGAGCGCACCATCTTCTTC AAGGACGACGGCAACTACAAGACCCGCGCCGAGGTGAAGTT CGAGGGCGACACCCTGGTGAACCGCATCGAGCTGAA | Template (as part of the plasmid pEGFPN1) for synthesis of polyFAM probes with 160 bp length |
| 2 | eGFP-PCR-F | [Bio]TTTTTTTTTTTTTTT-[Spacer C3]- CGCTACCCCGACCACATGAAG | Primers for PCR synthesis of polyFAM probes with 160 bp length |
| 3 | eGFP-PCR-R | [FAM]TTCAGCTCGATGCGGTTCAC | |
| | invA-PCR | CATGAAATGGCAGAACAGCGTCGTACTATTGAAAAGCTGTC TTAATTTAATATTAACAGGATACCTATAGTGCTGCTTTCTCTA CTTAACAGTGCTCGTTTACGACCTGAATTACTGATTCTGGTAC TAATGGTGATGATCATTTCTATGTTCGTCATTCCATTACCTACC TATCTGGTTGATTTCCTGATCGCACTGAATATCGTACTGGCGA TATTGGTGTTTATGGGGTCGTTCTACATTGACAGAATCCTCAG TTTTTCAACGTTTCCTGCGGTACTGTTAATTACCACGCTCTTTCG TCTGGCATTATCGATCAGTACCAGTCGTCTTATCTTGATTGAAG CCGATGCCGGTGAAATTATCGCCACGTTCGGGCAATTCGTTAT TGGCGATAGCCTGGCGGTGGGTTTTGTTGTCTTCTCTATTGTCA CCGTGGTCCAGTTTATCGTTATTACCAAAGGTTCAGAACGTGTC GCGGAAGTCGCGGCCCGATTTTCTCTGGATGGTATGCCCGGTA AACAGATGAGTATTGATGCCGATTTGAAGGCCGGTATTATTGA TGCGGATGCCGCGCGCGAACGGCGAAGCGTACTGGAAAGGGA AAGCCAGCTTTACGGTTCCTTTGACGGTGCGATGAAGTTTATCA AAGGTGACGCTATTGCCGGCATCATTATTATCTTTGTGAACTTTA TTGGCGGTATTTCGGTGGGGATGACTCGCCATGGTATGGATTTG TCCTCCGCCCTGTCTACTTATACCATGCTGACCATTGGTGATGGTC TTGTCGCCCAGATCCCCGCATTGTTGATTGCGATTAGTGCCGGTT TTATCGTGACCCGCGTAAATGGCGATACGGATAATATGGGGCG GAATATCATGACGCAGCTGTTGAACAACCCATTTGTATTGGTTGT TACGGCTATTTTGACCATTTCAATGGGAACTCTGCCGGGATT | DNA-target (invA gene fragment of *S.* Typhimurium) amplified by PCR |
| 4 | invA-PCR-F | CATGAAATGGCAGAACAGCG | Primers for PCR synthesis of dsDNA-target (fragment with 959 bp length of invA gene) of *Salmonella* Typhimurium |
| 5 | invA-PCR-R | AATCCCGGCAGAGTTCCC | |
| | invA-LAMP | GACCTGAATTACTGATTCTGGTACTAATGGTGATGATCATTTC TATGTTCGTCATTCCATTACCTACCTATCTGGTTGATTTCCTGA TCGCACTGAATATCGTACTGGCGATATTGGTGTTTATGGGGTC GTTCTACATTGACAGAATCCTCAGTTTTTCAACGTTTCCTGCGG TACTGTTAATTACCACGCTCTTTCGTCTGGCATTATCGATCAGT ACCAGTCGTCTTATCTTGATTGAAGCCGAT | DNA-target (invA gene fragment of *S.* Typhimurium) amplified by LAMP |

| | | | |
|---|---|---|---|
| 6 | invA-LAMP-F3 | GACCTGAATTACTGATTCTGGT | Primers for LAMP of *Salmonella* Typhimurium, designed by (Xia et al. 2021) |
| 7 | invA-LAMP-B3 | ATCGGCTTCAATCAAGATAAGA | |
| 8 | invA-LAMP-FIP | AACACCAATATCGCCAGTACGATC GTCATTCCATTACCTACCTA | |
| 9 | invA-LAMP-BIP | TTTCAACGTTTCCTGCGGTACTACT GGTACTGATCGATAATGC | |
| 10 | invA-LAMP-LF | TGCGATCAGGAAATCAACCA | |
| 11 | invA-LAMP-LB | ACCACGCTCTTTCGTCTG | |
| 12 | gRNA-invA1 | UAAUUUCUACUAAGUGUAGAUAAUACCGCCAAUAAAGUUCA* | Guide RNA recognizing dsDNA-target of *Salmonella* Typhimurium, designed by Luo et al. (2022) |
| 13 | gRNA-invA2 | UAAUUUCUACUAAGUGUAGAUCUGCGGUACUGUUAAUUACC* | Guide RNA recognizing dsDNA-target of *Salmonella* Typhimurium, designed by Xia et al. (2021) |
| 14 | ROX-dT15-BHQ2 | [ROX]TTTTTTTTTTTTTTT[BHQ2] | Conventional probe for CRISPR/Cas12a assay with fluorescence detection |
| 15 | FAM-dT15-Bio | [FAM]TTTTTTTTTTTTTTT[Bio] | Conventional probe for CRISPR/Cas12a assay with LFT detection |
| 16 | tripod-dT15 | ([FAM][HEg])3[TREBLER]TTTTTTTTTTTTTTT[Bio] | Proposed probes for CRISPR/Cas12a assay with LFT detection |
| 17 | tripod-dT50 | ([FAM][HEg])3[TREBLER]TTTTTTTTTTTTTTTTTTTT TTTTTTTTTTTTTTTTTTTTTTTTTTTTTT[Bio] | |

The abbreviations in the table mean [Bio] – bioitin, [FAM]—fluorescein, ROX – 5-carboxyrhodamine-X, BHQ2 – black hole quencher2, HEg – hexaethylene glycol, [TREBLER]—branch point in 5′-terminal direction due to using trebler phosphoramidite for synthesis of oligonucleotides; [Spacer C3] – propandiol.

* Site of recognition is denoted by the single or double underline

**Section 2.** Synthesis and characterization of gold nanoparticles (GNPs) and their conjugates with anti-fluorescein antibodies (antiFAM)

For synthesis of gold nanoparticles (GNP), one milliliter of 1% $HAuCl_4$ was added to 97.5 mL of deionized water. The mixture was continuously stirred and heated to the boiling point; then, 1.5 mL of 1% sodium citrate was added. The GNP solution was continuously boiled for another 30 min, then cooled and stored at 4 °C. The GNPs were dropped onto a grid (300 mesh, Pelco International, USA) coated with polyvinyl formal support film. The images were obtained with a CX-100 electron microscope (Jeol, Japan) at an accelerating voltage of 80 kV. The nanoparticles characteristics were obtained with Image Tool software (University of Texas Health Science Center in San Antonio, USA) (**Figure S1**).

The GNPs were conjugated with antibodies specific to fluorescein (antiFAM) through physical adsorption. The solution of GNP was adjusted to pH 9.0. Then 12 µg antiFAM was added to each 1 mL of GNP solution (1 nM). The synthesis was carried out at RT for 1 h, with continuous mixing. Afterwards, BSA was added to a final concentration equal to 0.25%. The mixture was centrifuged at 15,000× g for 30 min to separate the GNP conjugates. The synthesized GNP conjugates were resuspended in conjugate buffer (0.1M Tris-HCl (pH 8.6), 1% BSA, 1% trehalose, 0.1% polyvinylpyrrolidone 25, 0.1% Tween-20, 0.1% Na azide).

The spectra of the GNPs and their conjugates were recorded using a Libra S80 spectrophotometer (Biochrom, UK). The results are presented in **Figure S2a, b**. The hydrodynamic diameters of the GNPs and conjugates were measured using Zetasizer Nano (Malvern Panalytical, Malvern, UK). All measurements were performed at 25 °C, and scattering angle was equal to 173°. The results are presented in **Figure S2c,d**.

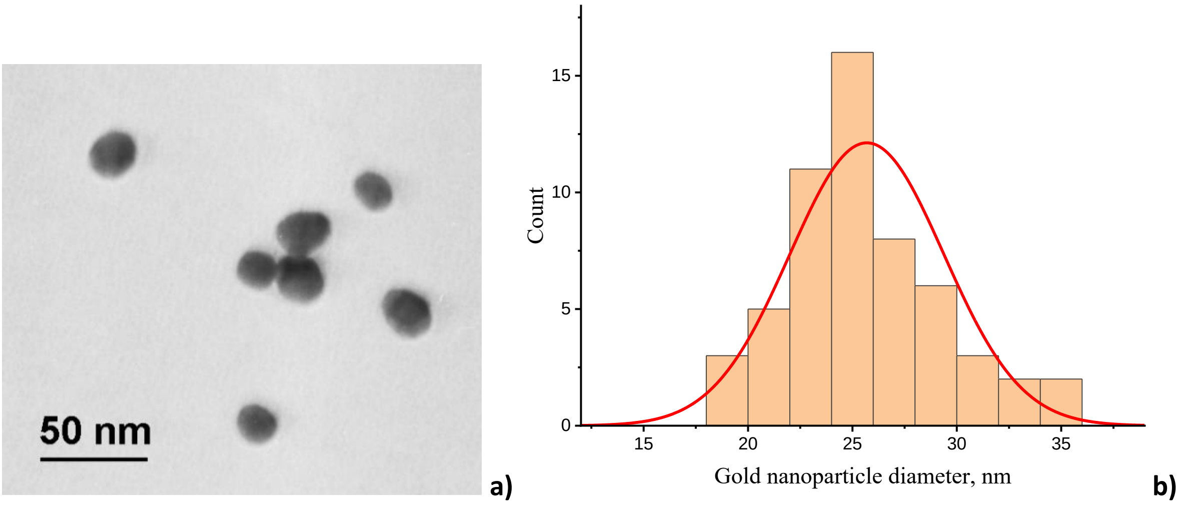

**Figure S1.** Transmission electron microscopy of GNPs: image of GNPs **(a)** and histogram of GNP diameter (n = 56) and fitting by Gauss approximation according OriginPro 9.0 software (Origin Lab, USA) **(b)**. The mean diameter is 25.7 nm, and the standard deviation is 3.7 nm, a degree of ellipticity is 1.2±0.1.

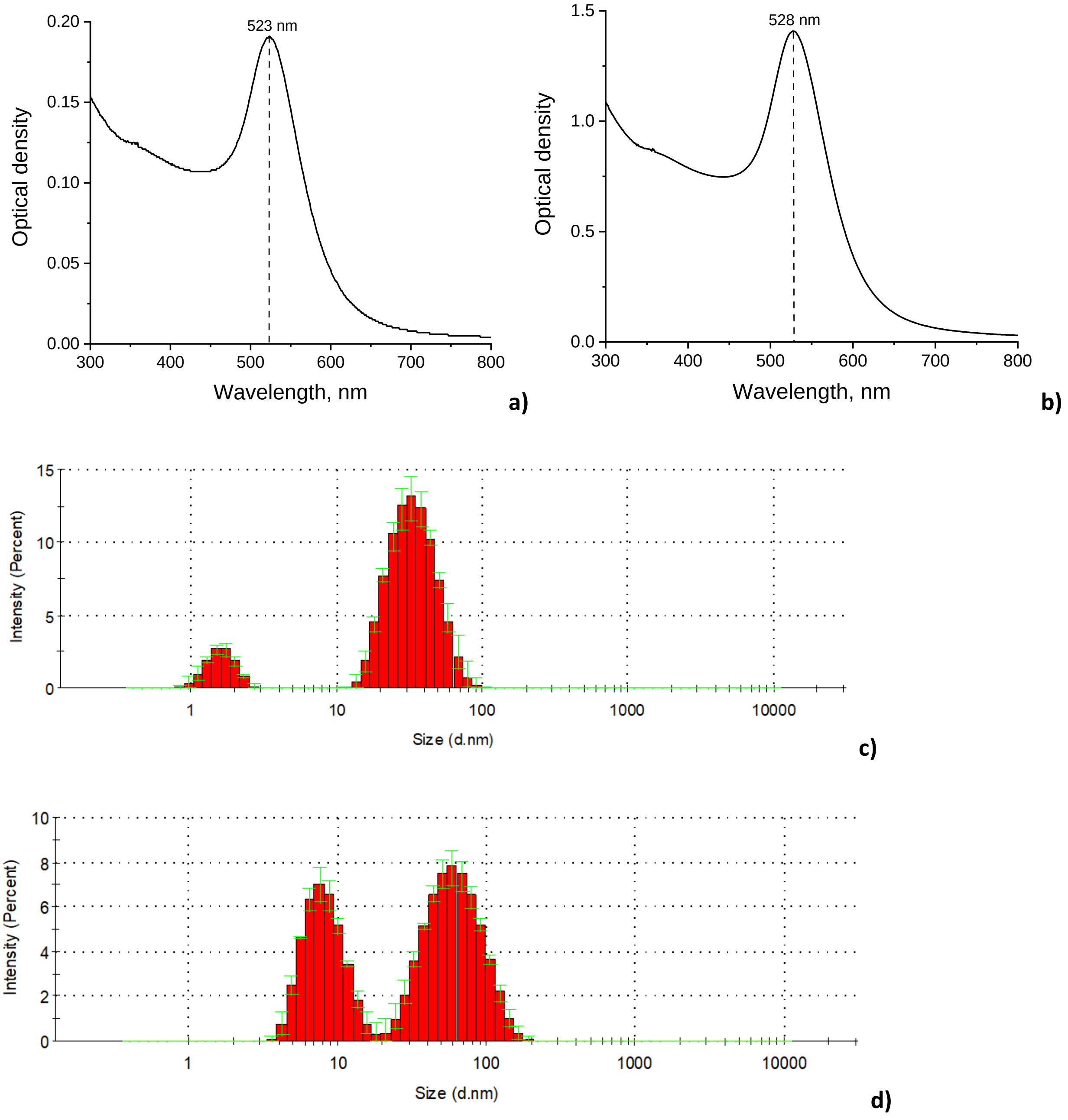

**Figure S2**. Absorption spectra of GNP **(a)** and antiFAM – GNP conjugate **(b)**. Distribution of hydrodynamic diameters of GNP, $D_H$ = 34.0 nm **(c)** and antiFAM – GNP conjugate, $D_H$ = 63.9 nm **(d).**

**Section 3.** Structure of trebler phosphoramidite using for synthesis of tripod probes

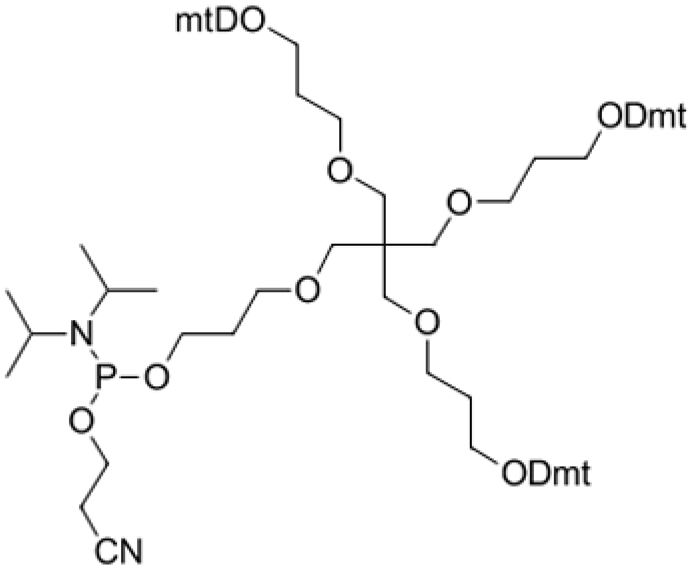

**Figure S3.** Structure of long trebler phosphoramidite ($C_{89}H_{107}N_2O_{15}P$, Mr = 1475.78 Da) using for synthesis of branched DNA structures where Dtm is dimethoxytrityl protecting group.

**Section 4.** Characteristics of polyFAM probes with LFT-common and LFT-CRISPR

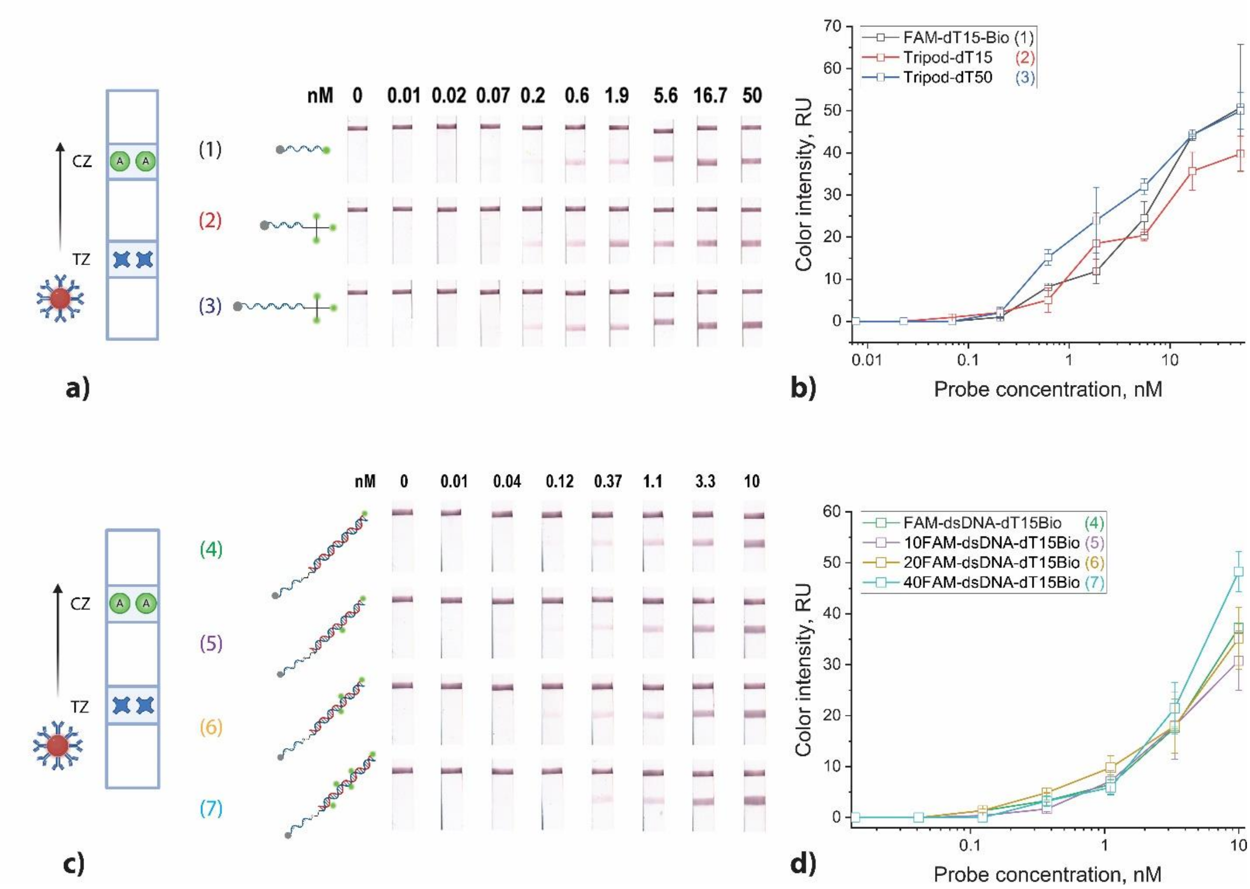

**Figure S4.** Results of LFT-common testing with immobilized streptavidin in the test zone, obtained for single-labeled probes: FAM-dT15-Bio **(a)**, **(b)** and FAM-dsDNA-dT15-Bio **(c)**, **(d)**, and polyFAM-labeled probes: tripod-dT15, tripod-dT50 **(a)**, **(b)**; 10FAM-dsDNA-dT15-Bio, 20FAM-dsDNA-dT15-Bio, 40FAM-dsDNA-dT15-Bio **(c)**, **(d)**, in the concentration range of 0.01 to 50 nM. Test strip scans **(a)**, **(c)**, and concentration dependencies obtained for the test zone **(b)**, **(d)**.

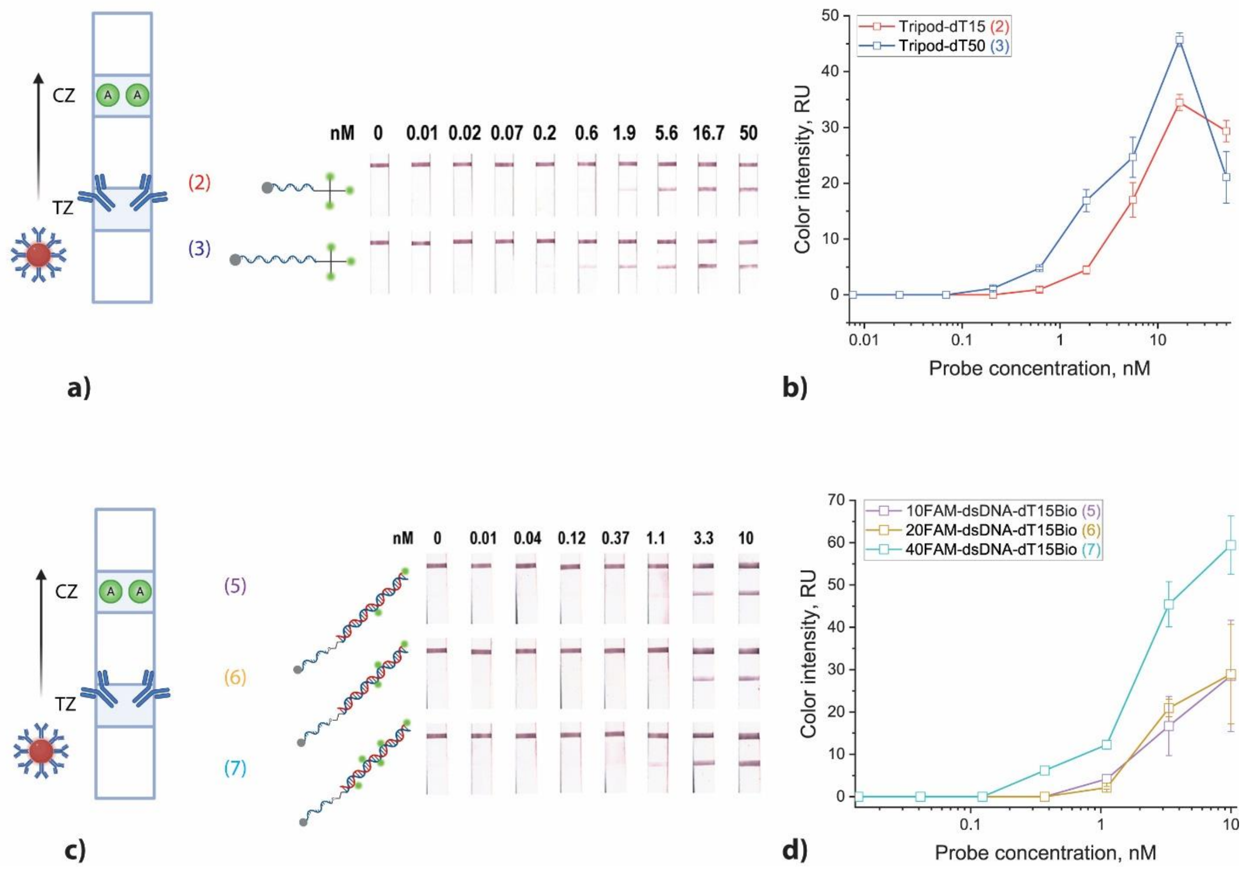

**Figure S5**. Results of LFT-CRISPR testing with immobilized antiFAM antibodies in the test zone, obtained for polyFAM-labeled probes: tripod-dT15, tripod-dT50 **(a)**, **(b)**; 10FAM-dsDNA-dT15-Bio, 20FAM-dsDNA-dT15-Bio, and 40FAM-dsDNA-dT15-Bio **(c), (d)** in the concentration range of 0.01 to 40-50 nM. Test strip scans **(a)**, **(c)**, and concentration dependencies obtained for the test zone **(b)**, **(d)**.

**Section 5.** Characterization of dsDNA target of *S.* Typhimurium obtained by PCR

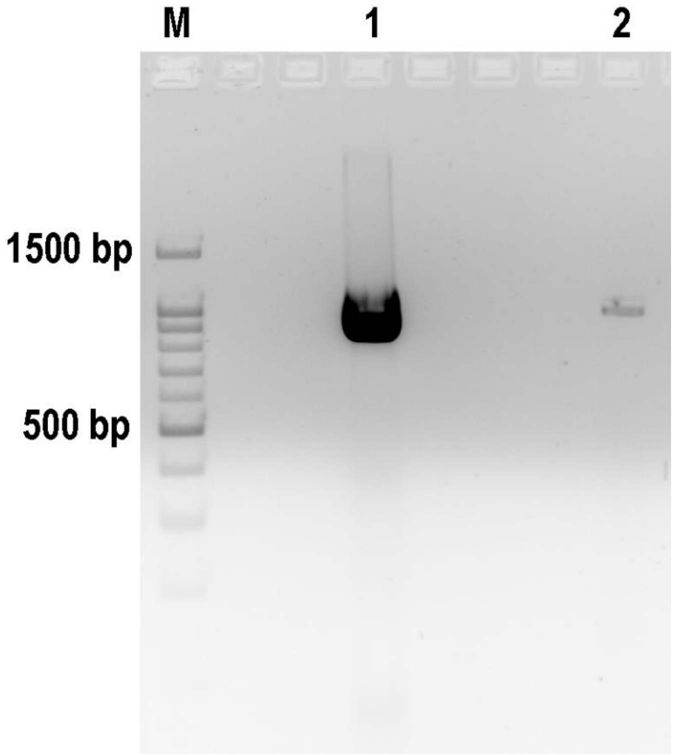

**Figure S6**. Visualization of PCR products (DNA fragment of invA gene with length of 959 bp) after electrophoresis in 2% agarose gel for the *S.* Typhimurium. M is the DNA marker, 1 is the PCR product after concentration, 2 is the diluted aliquot of PCR product after purification.

**Section 6.** Basic check of the CRISPR/Cas12a test system including tripod-dT15 and LFT-CRISPR

**Tables S2.** Testing tripod-dT15 probe using LFT-CRISPR

| № | gRNA | Cas12a | dSDNA target, 2 nM | tripod-dT15-probe, 33 nM | MP-Str | LFT-CRISPR | Mean color intensity of TZ ± sd (n = 2), RU |
|---|---|---|---|---|---|---|---|
| 1 | - | - | - | + | - | 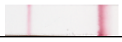 | 12.3 ± 0.7 |
| 2 | - | + | - | + | - | 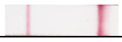 | 18.1 ± 3.0 |
| 3 | - | + | - | + | + | 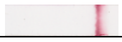 | 0 |
| 4 | + | + | - | + | + | 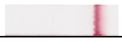 | 0 |
| 5 | + | + | + | + | + | 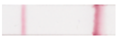 | 15.0 ± 0.6 |

**Section 7.** Results obtained with loop-mediated isothermal amplification (LAMP)

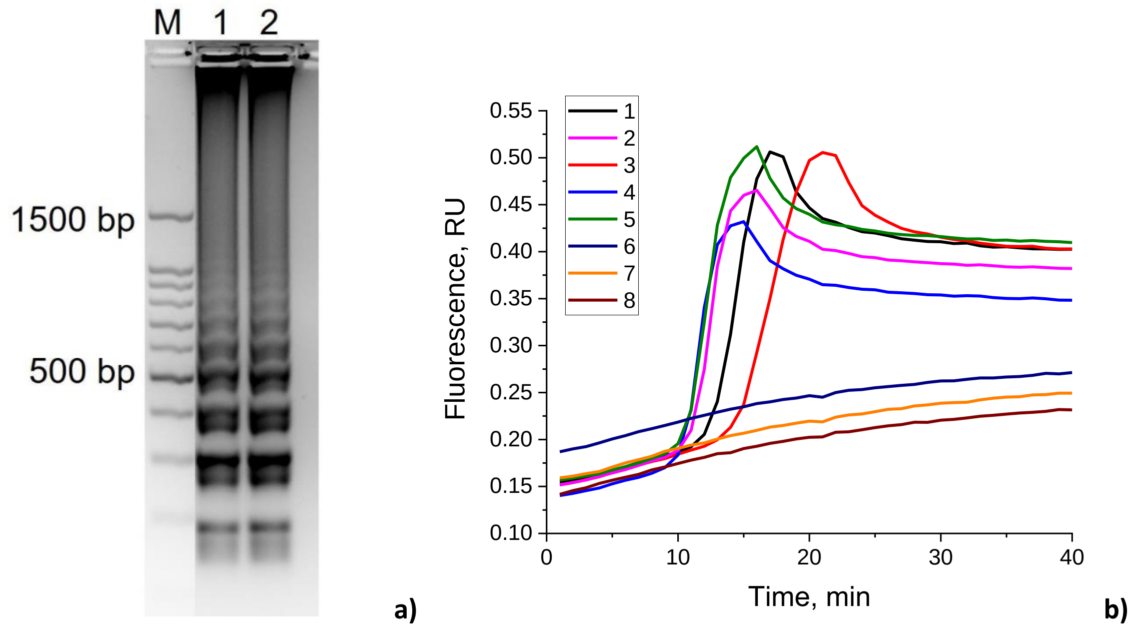

**Figure S7**. LAMP products visualized with 2% agarose gel electrophoresis stained with ethidium bromide **(a)** and SYBR Green I staining **(b)** for total DNA isolated from inactivated cells of *S.* Typhimurium (1), *S.* Enteritidis (2), *S.* Paratyphi (3), *S.* Virchow (4), *S.* Anatum (5), *Escherichia coli* 0157:H7 (6), *Yersinia enterocolitica* (7); TAE buffer (8).

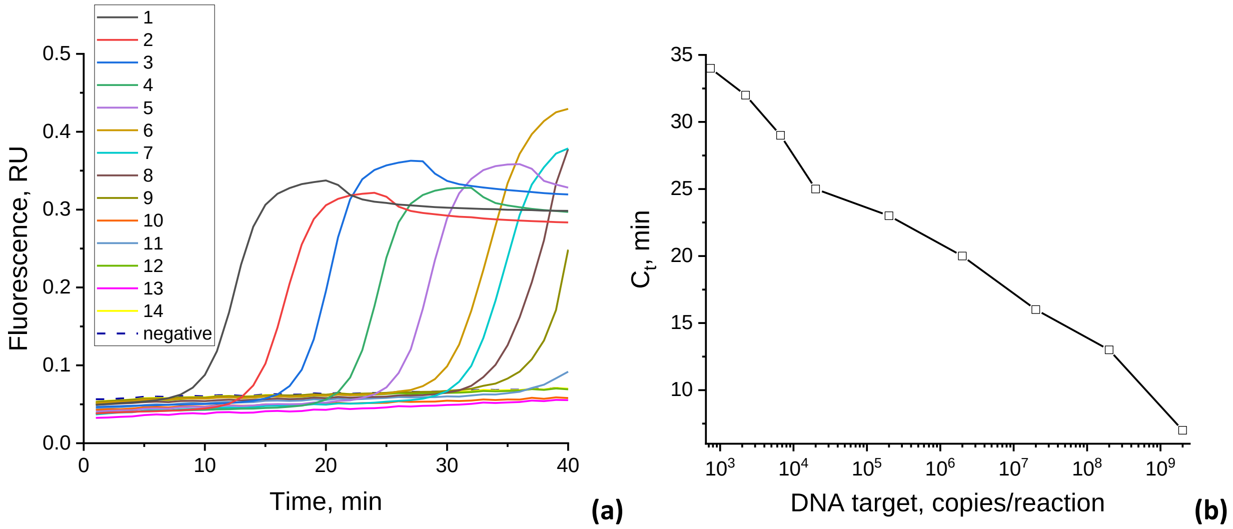

**Figure S8**. LAMP with SYBR Green I detection for DNA target (959 bp, invA gene of *S.* Typhimurium) at different concentrations: 2 × $10^9$ (1), 2 × $10^8$ (2), 2 × $10^7$ (3), 2 × $10^6$ (4), 2 × $10^5$ (5), 2 × $10^4$ (6), 6.6 × $10^3$ (7), 2.2 × $10^3$ (8), 7.4 × $10^2$ (9), 2.5 × $10^2$ (10), 82 (11), 27 (12), 9.1 (13), 3 (14) DNA copies per reaction, negative – TAE buffer. Fluorescence curves **(a)**, concentration dependence of signal appearance (Ct) on the concentration of DNA target **(b)**.

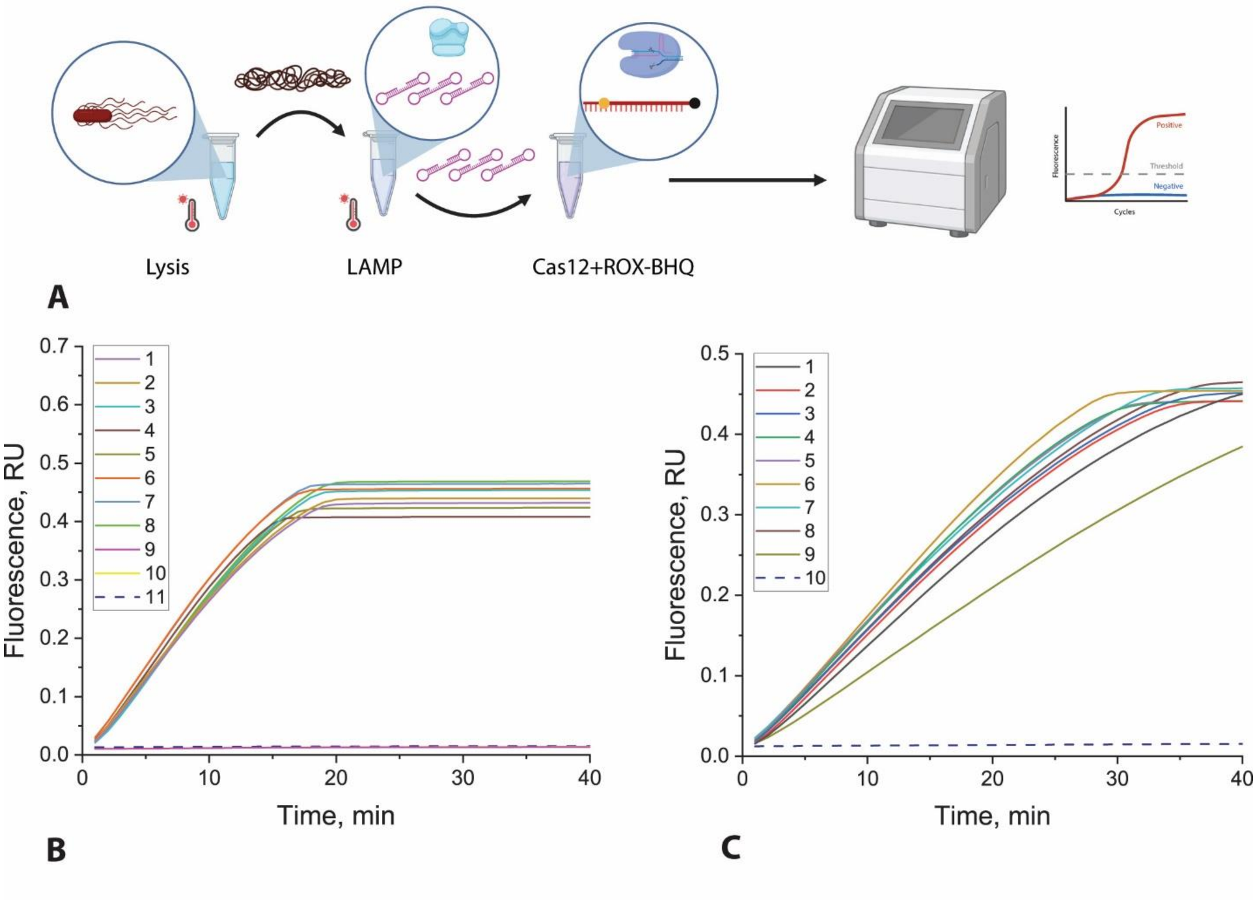

**Figure S9**. CRISPR/Cas12a for LAMP samples. Scheme of CRISPR/Cas12a with ROX-dT15-BHQ2 **(A),** fluorescence curves obtained for ROX-dT15-BHQ2
with LAMP of dsDNA (959 bp, invA gene of *S.* Typhimurium) **(B)**, where numbers indicate: 1 – 2 × $10^5$, 2 – 2 × $10^4$, 3 – 6.6 × $10^3$, 4 – 2.2 × $10^3$, 5 – 740, 6 – 250, 7 – 82, 8 – 27, 9 – 9, 10 – 3, 11 – 0 copies/reaction; with LAMP of *S.* Typhimurium cells after thermolysis **(C)**, where numbers indicate: 1 – 2 × $10^4$, 2 – 2 × $10^3$, 3 – 2 × $10^2$, 4 – 67, 5 – 22, 6 – 7, 7 – 3, 8 – 1, 9 – 0.3, 10 – 0 cells/reaction.

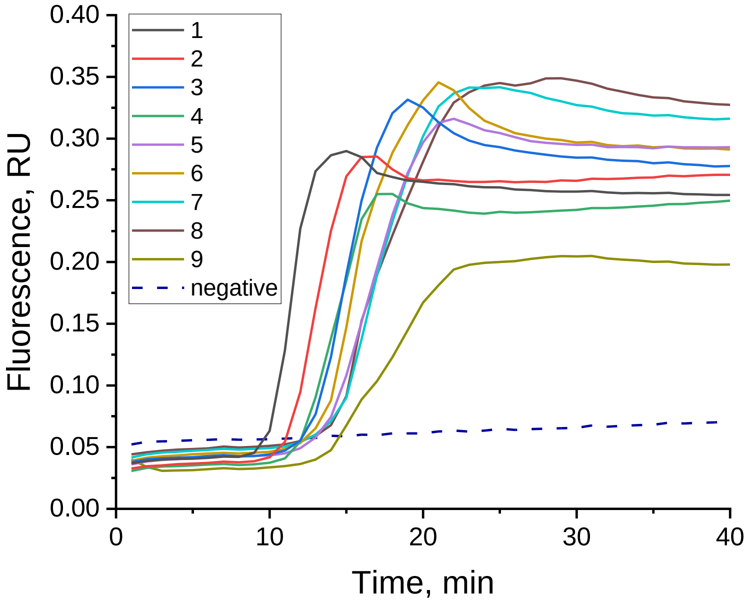

**Figure S10.** Fluorescence curves obtained by LAMP with SYBR Green I detection for *S.* Typhimurium cells in buffer after thermolysis, where numbers indicate: 1 – 2 × $10^4$, 2 – 2 × $10^3$, 3 – 2 × $10^2$, 4 – 67, 5 – 22, 6 – 7, 7 – 3, 8 – 1, 9 – 0.3 cells/reaction, negative – TAE buffer.

**Section 8.** Comparison of CRISPR/Cas12a test-system combined with LFT for *Salmonella* detection

**Table S3.** The comparison of CRISPR/Cas12a test-system combined with LFT for *Salmonella* detection

| **Amplification** | **Type of LFT-detection** | **Label in LFT** | **Detection limit** | **Reference** |
|---|---|---|---|---|
| RPA*-CRISPR/Cas12a | competitive | gold nanoparticle | 384 CFU/mL | (Chen et al.) |
| LAMP–CRISPR/Cas12a | competitive | gold nanoparticle | 1.22 CFU/mL | (Lee and Oh 2023) |
| LAMP–CRISPR/Cas12a with bifunctional nanoflower nanozyme | sandwich | gold nanoparticle | $10^2$ CFU/mL | (Qiu et al. 2024) |
| RAA**-CRISPR/Cas12a | DETECTR | gold nanoparticle | 33 CFU/mL | (Wang et al. 2024) |
| RPA-CRISPR/Cas12a | competitive | fluorescent label | 1 copy/µL | (Yuan et al. 2024) |
| LAMP-CRIPR/Cas12a | sandwich | gold nanoparticle | 150 cells/mL | This study |

*RPA – recombinase polymerase amplification, **RAA – recombinase-assisted amplification
